# A novel approach to quantifying inter-individual distance in insects

**DOI:** 10.1101/2022.06.28.497925

**Authors:** Kristina Corthals, Lennart Hüper, Sven Neumann, Ralf Heinrich, Heribert Gras, Bart R.H. Geurten

**Affiliations:** Lund University, Department of Experimental Medical Science, Sölvegatan 35, 221 00 Lund; Georg-August-University Göttingen, Department of Cellular Neuroscience, Julia-Lermontowa-Weg 3, 37077 Göttingen; Georg-August-University Göttingen, Feinmechanische Werkstätten, Julia-Lermontowa-Weg 3, 37077 Göttingen

**Keywords:** *Drosophila*, insect, courtship, social behaviour, aggression, sexual dimorphism, group behaviour

## Abstract

1

Behaviour involving two or more individuals within the same species is known as social behaviour. Sexual dimorphisms and environmental cues as well as age, experience and social hierarchy shape social behaviour. The study of this complex behaviour, however, has one essential boundary condition: the distance between individuals. Because social signals (e.g. acoustic, visual or olfactory) have limited reach the inter-individual distance is crucial for the correct transmission of such signals. In this study we present a newly developed assay to study group behaviour and inter-individual distance in *Drosophila melanogaster*. This assay consists of a circular observation space flanked by two automatic release devices that allow flies to enter individually. By releasing the flies one at a time, the observer can study different features of (intra-)group behaviour with different group compositions. Importantly, the observer can control (manually or through automatisation) the increase of group size over time.

Over the last decades insects became more relevant as convenient model organisms to study the neurophysiological and genetic basis of (human) neuro-developmental disorders. *Drosophila* models are used to study the genetic and neuronal underpinnings of a wide range of neurological disorders. In some cases the studies revealed alterations in social behaviour consistent with descriptions of behavioural symptoms in human disorders.

Social behaviours in *Drosophila* are well-studied and include courtship, mating, aggression and group interactions. This setup will facilitate the analysis of these aspects of group interactions in *Drosophila*, allowing for a deeper understanding of the neuronal circuits and genetic factors involved in those behaviours.

**Contribution to the field:** Social behaviour pertains to the most sophisticated behavioural feats, as it involves multiple, interacting individuals. These complex interactions often conceal the underlying neuronal and ethological mechanisms. One of the most basal ethological mechanisms is the inter-individual distance, which resembles a perimeter in which each individual needs to formulate a response to the approach of others.

We introduce a device that allows to test the inter-individual distance under consistent circumstances, by automating the entry time and direction of conspecifics into the arena. We can further observe the composition, dynamics and forming of larger animal groups as well as their separating. We can also observe how the individual distances alter during the process. Also other behaviours can be easily observed, e.g. aggression, courtship, homosexual courtship, etc. We successfully employed this approach in [1] and could discriminate the role of different neuroligins in social behaviour. We provide a detailed description including building plans and material lists for this social observation device. The system can be run in an automatic mode to ensure the consistency of experiments or in a manual mode to test animals under more flexible social situations. We provide multiple back lighting systems to test animals in the dark (infra-red LEDs) or in illuminated environments (vis. range LEDs). The system is fully automated and can be linked with a number of animal trackers (e.g. T-Rex, deeplabcut, LACE, etc.) via simple videography. We hope that our experimental setup augments the variety of behaviours testable in ethological setups (T-maze, water-mazes, operant conditing setups, etc) with social interaction and group formation.

## 4 Introduction

The observation of social behaviour is intrinsically complicated as in contrast to other ethological topics the stimuli in social contexts are other animals and their flexible behaviour [2]. This makes it nearly impossible to create consistently identical experimental situations for the observed animal. In order to minimise other sources of random stimuli we tried to create a setup that guides the approach of individuals to each other as consistent and as reproducible as possible. Therefore, we generated a device that allows the experimenter to control direction and timing of the approach of two individuals and the forming of groups. The formation of groups depends on a variety of factors: genetic relationships can favour the formation of long-lasting groups but might also be limited by the duration of nursing or breeding [3, 4]. Group formation can also be found in a range of cumulative behaviours such as swarming in locusts [5, 6], nest building in ants [7, 8], migration in flocks of birds [9–11], courtship and mating in garter snake males [12], collective decision making in schools of fish [13, 14] and the cellular slime mould *Physarum polycephalum* [15].

The mechanics of group interactions, however, remain mostly elusive, as these behaviours are influenced by multiple environmental and internal factors. Both mammals and insects have distinct groups of neurons regulating social interactions like mating and aggression [16, 17]. Additionally, neuromodulators like tachykinin, serotonin and oxytocin as well as species-specific pheromones are controlling a wide range of social behaviours [16, 18, 19].

In *Drosophila* communication with con-specifics is mediated by auditory, olfactoy, visual and gustatory cues [19–21]. Auditory cues include courtship sounds produced by males at a distance of 1 mm to 4 mm from the female [22].

Here, we introduce a behavioural setup that targets the most basic characteristic of group interactions: the inter-individual distance. The inter-individual distance marks the defence parameter of each animal. To shorten the inter-individual distances most species developed a particular strategy to approach con-specifis to avoid aggressive interaction. For example *Drosophila* developed a complex mating behaviour that starts with long range acoustic communication [23] and continues through a series of prototypical behaviours that shortens the inter-individual distance until copulation is reached [19, 24, 25]. These mating behaviours depend on olfactory, visual, auditory or gustatory communication cues to be initiated and each of these signals has a certain limit of reach. Therefore, the distance between the individuals determines which behaviours and communication signals are possible. Inter-individual distance is further influenced by other factors like age, hierarchy, previous experiences, available space or neuronal alterations. In *Drosophila* deficiency of the postsynaptic adhesion molecules Drosophila neuroligin 2 and 4 (Dnlg2 and Dnlg4) leads to disrupted group formation and an increase in inter-individual distance [1], consistent with previous reports of abnormal courtship and social behaviours in Dnlg2-deficient *Drosophila* [26]. Neuroligins are well-described post-synaptic proteins involved in the formation and maturation of synapses [27–31]. Neuroligins are present in both mammals and insects and recent studies show their conserved function mediating social interaction and communciation [1, 26, 32–34]. In humans, impairment of neuroligins have been associated with ASDs and alterations in social behaviour and communication [35–38].

Cooperation often leads to immediate benefits for all members of the cooperative group. Cooperation and its benefits rely on genetic relationships, reciprocity, or both. Living in cooperative groups can offer several advantages over living in isolation: better protection against predators [39], facilitated foraging or hunting [40], and nest building [8], or more successful mating approaches [12, 41]. Furthermore, crucial behaviours such as mating, predator avoidance and foraging are learned by insects in a social context. Naïve bumblebees learn foraging choices by monitoring which flowers are the preferred targets from other, more experienced bumblebees [42–44]. *Drosophila* females are influenced in their mate choice by observing successfull and unsuccesfull copulation attempts of their conspecifics [45]. When faced with a predator, crickets learn and adjust their predator-avoidance behaviour based on more experienced demonstrators [46]. These examples show the importance of social interactions even in non-colonial animals.

Studies of group behaviours in the past were often limited to observations in the field. This involves catching and marking individual animals [47, 48] or using DNA fingerprinting to identify individuals and their relationship [49, 50]. More recent studies are using virtual reality (VR) technologies to simulate environments with several animals while assessing the behaviour of an idividual [51, 52]. These setups allow the control of several environmental factors and can imitate the presence of con-specifics using the distinct movement patterns of each species [52]. While these setups open up an impressive range of possible applications, they are technically very elaborate and often expensive. Behavioural setups that can be produced at low cost are important for small labs and education purposes [53]. Our new approach to quantify inter-individual distance in *Drosophila* will therefore be a convenient and easily accessible tool to unveil the basic mechanisms of forming and maintaining of groups. The design allows the experimenter to manually control several features of group formation, such as number of individuals, the increase of the group over time and group composition. Furthermore, it can be used to screen for alterations in social behaviours and communication in (new) *Drosophila* models for several neuro-developmental disorders.

As a proof-of-concept we are showing and discussing data on the inter-individual distance (IID) comparing wt *Drosophila* with a neuroligin knock-out strain.

## 5 Methods & Material

### 5.1 Animal rearing

All *Drosophila* strains were raised at 25 ^°^C 60% relative humidity at a 12:12 h dark/light cycle on standard apple juice medium (1 l apple juice, 500g fresh yeast, 500g sugar, 60g agarose, 20g salt, 250g flour and 30ml proprionic acid, water was added to an end volume of 7l). Canton-S was used as a wild-type control (obtained from Bloomington Drosophila Stock Center, #64349), dnlg-mutant strain was *dnlg2*^*ko17*^ (provided by Wei Xie, Southeast University, Nanjing, China; characterized by Corthals et al. 2017 [1]).

### 5.2 Setup and experimental procedure

This behavioural setup allows the investigation of group formation and interaction while manually controlling several features of the group without any manual animal handling.

The setup consists of a polyamide (PA) observation arena with a diameter of 60 mm a height of 2 mm, closed with lid made off anti-glare acrylic glass (see Figure 1). This prevents the flies from flying in the arena but allows them to freely walk without being manipulated beforehand (i.e. wings cut off or glued to the sides). The anti-glare acrylic glass is coated with SigmaCote (Sigma-Aldrich, SL2-100ML, St. Louis, Missouri, United States) before every trial to prevent the flies from walking on the ceiling, as this can interfere with the video tracking.

**Figure 1:**
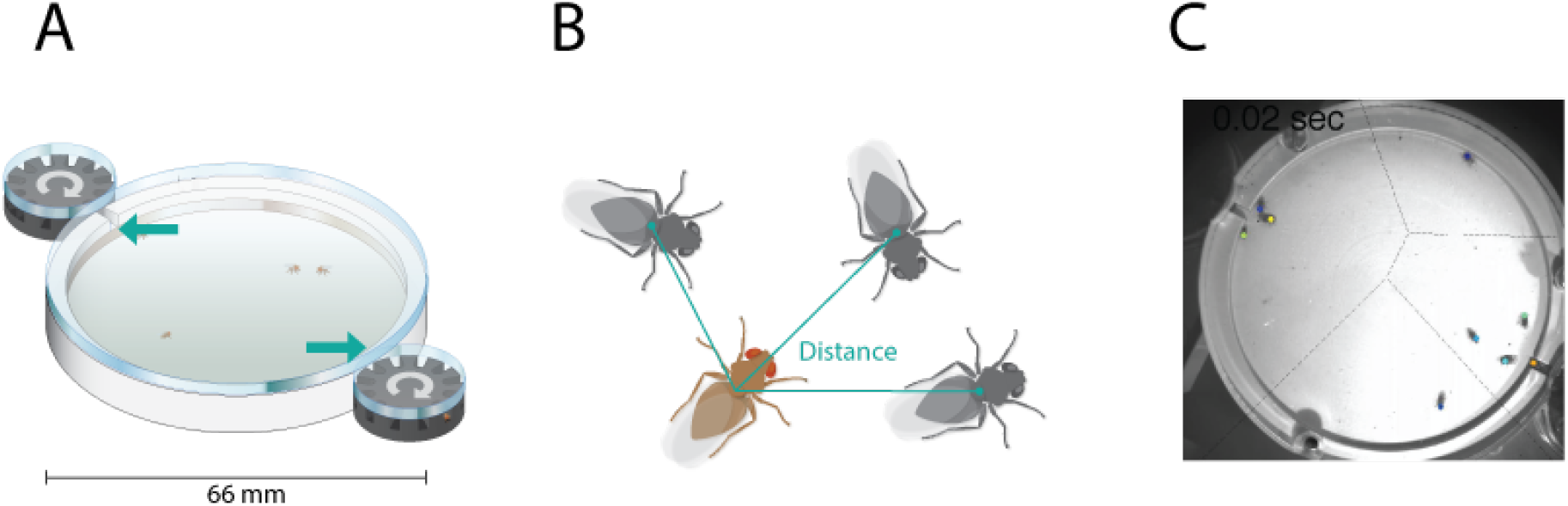
Concept drawing of the ethological setup. A) Concept drawing depicting the main observation chamber (white) and the two flanking entry revolvers (black). Both revolvers are connected via a small tunnel to the observation chamber. The tunnels are highlighted by COLOUR arrows. All three objects are covered by an acrylic glass pane. B) Euclidean distance between focus animal and the other individuals inside the arena. The distance is measured as the norm of the vector between two individuals. C) Example footage from the observational chamber. Colour dots represent tracked individuals. The dashed lines represent the borders of Voronoi cells [54] between groups based on hierarchical agglomerative clustering [55].

The arena is flanked by two revolvers, which are connected to the arena by a tunnel 3mm in lenght and can hold 12 seperated flies each, housing a maximum of 24 flies in a single experiment. Each revolver is connected to a 24-step-motor. These motors rotate a full 360^°^ in 24 evenly spaced steps, each covering a span of 15^°^. Each step of the motor either connects a fly to the tunnel leading into the observation arena or closes the tunnel connection. By flicking a switch the experimenter can rotate the revolver clockwise or anti-clockwise. If the flies should enter the arena at a scheduled time the experimenter can activate the automatic mode. The release period is set by a binary-coded-decimal (BCD) dip switch (ranging from 10 to 90 sec; SMC-D-111-AK-2, Hartmann-Electronic, Germany). During each period, one of the revolvers will open a fly enclosure to the tunnel, while the other one is set to closed position.

The motor-control-board is informed about the revolvers position by the zero point switch. This zero point switch responds to the 3mm notch in the driver socket (part #5; see p. 21) of the arena. The switch is mounted to the switch bracket (part #9; see p. 22) in a way that the lever points to the driver socket. When the notch passes the switch, the lever will fall into the notch and the motor control board registers that the switch is not longer pressed. Hence, for scheduled releases the board is able to detect the zero point position and than start releasing the flies based on a timer.

### 5.3 Electronic Control Circuits

The motors, lights, and connections are controlled via the central circuit board (see 31). An Arduino Micro (AT- mega32U4) functions as the central integrated circuit (IC) on the board. It receives inputs through the switches and buttons on the front and controls the two motors (RDM51/6, Berger Lahr, Austria) via half-H motor-drivers (L293D, Texas Instruments, USA) accordingly. There are also TTL output and input lines, so that the box can be triggered by video recording software or it can trigger the start of recording itself. A micro-USB port allows to change the programming of the Arduino later on. The R-78B5.0-1.5 (Recom Power, USA) supplies power to the system including motors and lights (see 31 ff.) and is connected to the current net via a 3A, 12V power supply. All cables are gathered in a D-sub connector, which connects it to the arena.

### 5.4 Embedded Code - Arduino

The Arduino runs a custom made code which is available at https://github.com/zerotonin/socialObservationStepMotorControl and based on the stepper motor control library Stepper available at https://www.arduino.cc/en/reference/stepper. The code consists of three scripts: (i) the main script that checks for button presses on the control device, runs motor initialisation, manual control of the motor, and starts experiments; (ii) the motor initialisation script includes a zero point calibration and can be found in the INIT_MOTORS.ino; (iii) the automation script for experiments controlled by a timer is written down in AUTOMATE.ino. Further documentation is given on the repository at github.com

### 5.5 Building Instructions

The motorised arena consists of thirteen different parts which are listed on the component part list (p. 18), including the needed quantity and the base material. Most parts can be produced with either a CNC-mill or a 3D printer. Parts 10-13 might be best manufactured using a lathe or 3D printer. You can see the assembly of the different parts in the so called assembly overview on page 19.

**Step 1**: The first step in the assembly is to position the step motors in the motor bracket (p. 20). Each motor bracket has thread-holes arranged at 120 ^°^. By inserting a grub screw into them, the motor can be fixed into the socket. The cables from the motors should be let out by the 14mm furrow in the motor bracket.

**Step 2**: Screw the electric switch to switch bracket (p. 22) and fix it to the motor bracket on the opposite side of the 14 mm furrow.

**Step 3**: Insert the cylindrical pins (component #10) into the driver-socket (p. 21) and install it on top of the stepper-motor. Fix the driver socket to the motor with another grub-screw.

**Step 4**: Now the base-plate (p. 23) can be installed on both motor contraptions and fixed with 8 M4×12 screws (component #12). To assure that the lids are always in place 4 pins (component #11) should be glued into the slots on the base-plate.

**Step 5**: The fly-revolver (p. 24) base can now be placed on top of the driver socket (p. 21). The revolver-lid (p. 25) can be placed above the revolver using the pins to guide it. Optionally one can insert a screw through the lid into the bottom of the revolver for easy handling. We strongly recommend this, but do not tighten it, otherwise the revolver will not turn or scratch the lid.

**Step 6**: Insert either the infrared LED panel or white LED panel into the trove in the base-plate. The white PA of the arena lets infrared and visible light travel through, allowing the observer to monitor the flies’ behaviour in perceived light or darkness. We used Nichia LEDs for white light (NSSW157AT-H3 SMD-LED, 27lm, 5000K, CRI 80) and 940 nm infra-red LEDs from Harvatek (HT-F157IRAJ IR emitter 940 nm 30 ^°^ 1206 SMD).

**Step 7**: The arena-lid (p. 30) can be screwed to the arena-plate (p. 29) and both should be inserted into the base-plate.

**Step 8**: Guide the step-motor wires and, if applicable, the LED wires out of the setup. We preferred to solder all wires to a D-sub connector as can be seen in supplementary material construction-photographs and 3D construction pdf.

### 5.6 Animal handling

To prepare the flies for the experiment they were collected in plastic tubes and immobilised by being kept on ice for 5 to 10 min. Contrary of using CO_2_ as an anaesthetic, chilling flies on ice has no prolonged effects on behaviour and is therefore the method of choice for behavioural experiments [56].

Upon reaching their chill coma, flies are tipped out of the vial onto an even metal surface submerged in ice to prevent them from waking up prematurely. Using blunt forceps, flies are subsequently loaded into the chambers of the revolver with their head facing outwards, to facilitate their passage into the arena. For the experiments shown here we used 12 males and females, respectively. Flies are given 15 min to fully recover from the chill coma before the experiment is started. Experiments were performed at 20 ^°^C in a temperature controlled room. The setup is placed in a box that can be closed to avoid disturbances during the experiment.

After starting the experiment the step motors below the revolvers transfer individual flies in front of the open tunnel. The flies than enter the arena within the desired interval chosen for the experiment. As the revolvers are dark (and very small) flies readily enter into the illuminated arena. The experiment ends when all flies have been released into the arena and group formation has been observed for a desired amount of time.

### 5.7 Tracking and data analysis

Flies were recorded at 50 fps using a Motion Traveller500 camera (IS, Imaging Solutions GmbH, Eningen, Germany) and TroubelPix video capturing software (NorPix Inc., Montreal, QC, Canada). Subsequently, videos were tracked using ivTrace (part of ivTools https://opensource.cit-ec.de/projects/ivtools by Dr. Jens P. Lindemann; Bielefeld University) and LACE [57]. Both trackers use image differences to track the centre of mass over the time course of the experiment, determining the flies position and orientation in a 2-dimensional space. There were no qualitative differences in tracking with both systems.

To avoid identity swaps between flies we calculated the squared euclidean distances between fly positions at frame f_n_ and fly positions at frame f_n+1_. These distances were used by an Hungarian algorithm [58] to sort flies according to their positions and verify the coherence of the data. Subsequently, the positional data was transformed from pixels to mm.

To assess inter-individual distances, the relative positions between all animals were calculated for the first frame of every second of the recording. To determine the position of one animal (A_*r*_) relative to the position of our focus animal (A_*O*_), we subtracted the position vector of the focus animal A_*O*_ from the position vector of A_*r*_. Hence we get a Cartesian vector pointing from our focus animals’ position to the observed animals’ position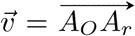. To account for the orientation of our focus animal we need to counter-rotate the vector 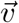 by the orientation angle of the focus animal. This way all focus animals are orientated similarly and can be compared. The rotation is performed by multiplying 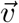 with the rotation matrix R, where *α* is the orientation angle of the focus animal:

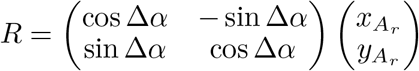

After this transformation, we convert 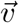 from a Cartesian to a polar coordinate system, consisting of a radius *r* and an angle *θ*, using the following equations:

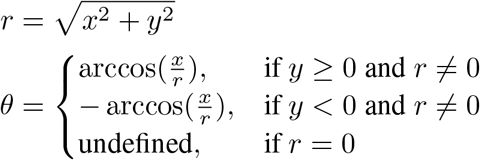

Subsequently, the relative positions of the animals were plotted as 2-dimensional histograms and the probability densities were calculated with a Gaussian kernel density estimation (KDE). Finally, the probability density of each histogram was normalised by bin surface area and then normalised so that the integral of the probability density was equal to 1.

## 6 Results

In this study we introduce a new behavioural setup that facilitates assessment of several aspects of group behaviour. It allows the monitoring of group formation and the quantification of multiple aspects of social behaviour, such as courtship, aggression or dominance behaviour [1]. As a proof of concept, we present data on the inter-individual distance (IID). The IID was quantified by measuring the euclidean distance between one fly, which we call the focus fly and all other individuals in the arena (Figure 1B).

Based on the IIDs, which can be derived from the individual fly positions (coloured dots in Figure 1C), we can amalgamate groups from individuals. We grouped individuals using machine learning algorithms to avoid observer bias [1]. A simple agglomerative cluster algorithm segregates the flies in Figure 1 C into 4 groups. Two groups consist of single flies, the other two groups are formed by three and four flies, respectively. Analysing the IIDs in a group revealed that the typical IID in groups is ca. 6 mm [1].

Augmenting the pure IID analysis, our setup also allows to analyse the orientation of the observed animal to the focus animal. We compare the IID of a wildtype *Drosophila* strain *CantonS* with a dnlg mutant strain *dnlg2*^*ko17*^ that has previously been shown to display impairment in social behaviours [1, 26]. In the wt group conspecifics are either located at a distance of about 10 mm and to the side or are in very close proximity to the focus individual and positioned either in front or at the back (Figure 2A). In comparison the visualized IDDs of the dnlg mutant line *dnlg2*^*ko17*^ show no clear preferred distribution, conspecifics are found at any distance and in any orientation towards the focus fly (Figure 2B). The IID to male or female flies differs in the wildtype strain. While distant females are predominantly focused to the sides, females closer to the focus animal are positioned to the front or the back (Figure 2C). Male *Drosophila* generate courtship song by extension and vibration of one wing which in turn activates female receptivity, producing two types of pulse song and one type of sine song [22, 59, 60]. The function of the sine song, intertwined with the interpulse interval, communicates the *Drosophila* species, to ensure mating compatibility [59, 60]. The pulse song are used to arouse the female and seperate in two types: P_fast_ and P_slow_ [22, 59]. P_slow_ is used for close range courtship, while P_fast_ is louder and used for far distance courtship. During the far distance stage of courtship males position their wing, producing P_fast_ song, towards the female antenna, the hearing organ of *Drosophila*, optimizing its reception [59, 61–63] Close distance courtship entails tapping the abdomen of the female as well as mounting to intiate courtship, which needs the female to be positioned either to the front or the back [64–66].

**Figure 2:**
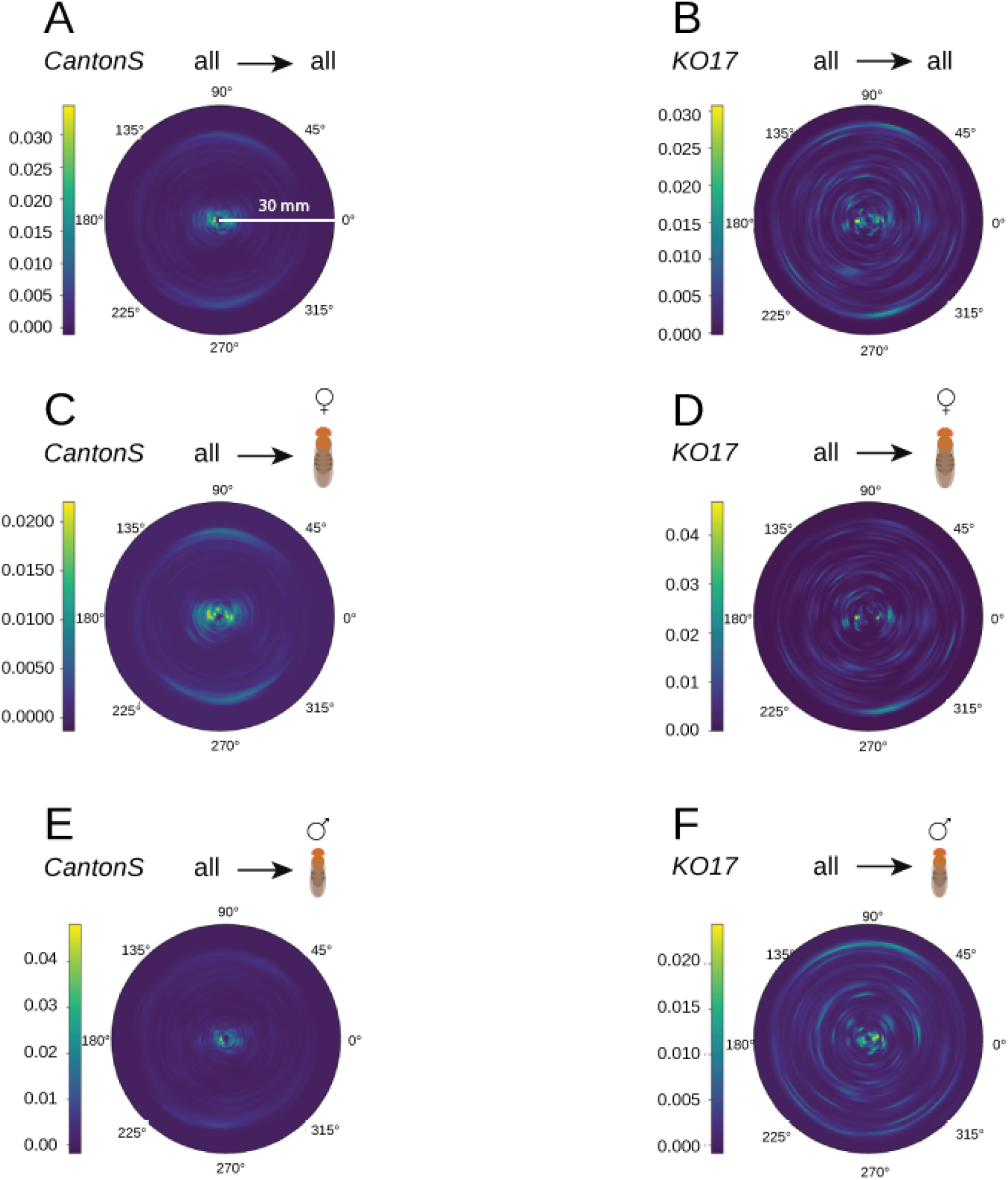
Histograms of inter-individual distances (IID) between flies. Each plot shows the colour coded probability distribution of fly positions around the focus animal. The colour code is given by the color bar left to the polar plot depicting the probability distribution. The head of the focus animal is oriented to zero degrees. 90^°^ is to the focus animal’s left. 180^°^ is directly behind the focus animal and 270^°^ is to the focus animal’s right. The white axis denotes the distance in mm. Above each polar plot the focus animal group and the conspecific group is depicted. A) shows the IIDs between all CantonS wildtype flies. B) shows the IID between all *dnlg2*^*ko17*^ mutant flies. C and D depict the respective subset of IIDs to observed female flies. E and F show the respective subset of IIDs to observed male flies.

In contrast, males at any distance are positioned uniformly around the focus animal (Figure 2E). If we perform the same analysis on *dnlg2*^*ko17*^ flies, we only find remnants of this behavioural sex dimorphism (Figure 2D-F). The distribution of conspecifics appears random and no clear pattern can be observed.

We further separated our data by the sex of the focus fly and the sex of the surrounding conspecifics. This revealed a clear sexual dimorphism in the occurring IIDs. Wt female focus flies keep their female conspecifics either at a very close distance range < 5 mm or at a distance of about 10 mm (Figure 3A). While *Drosophila* females show no agonistic behaviour towards other females in a courtship setting, female to female aggression behaviour has been reported in a context of sparse nutrient or competing for egg-laying sites [67, 68]. Since our data set includes both virgin and mated females we can assume that female-female aggression is indeed occurring in our setup.

**Figure 3:**
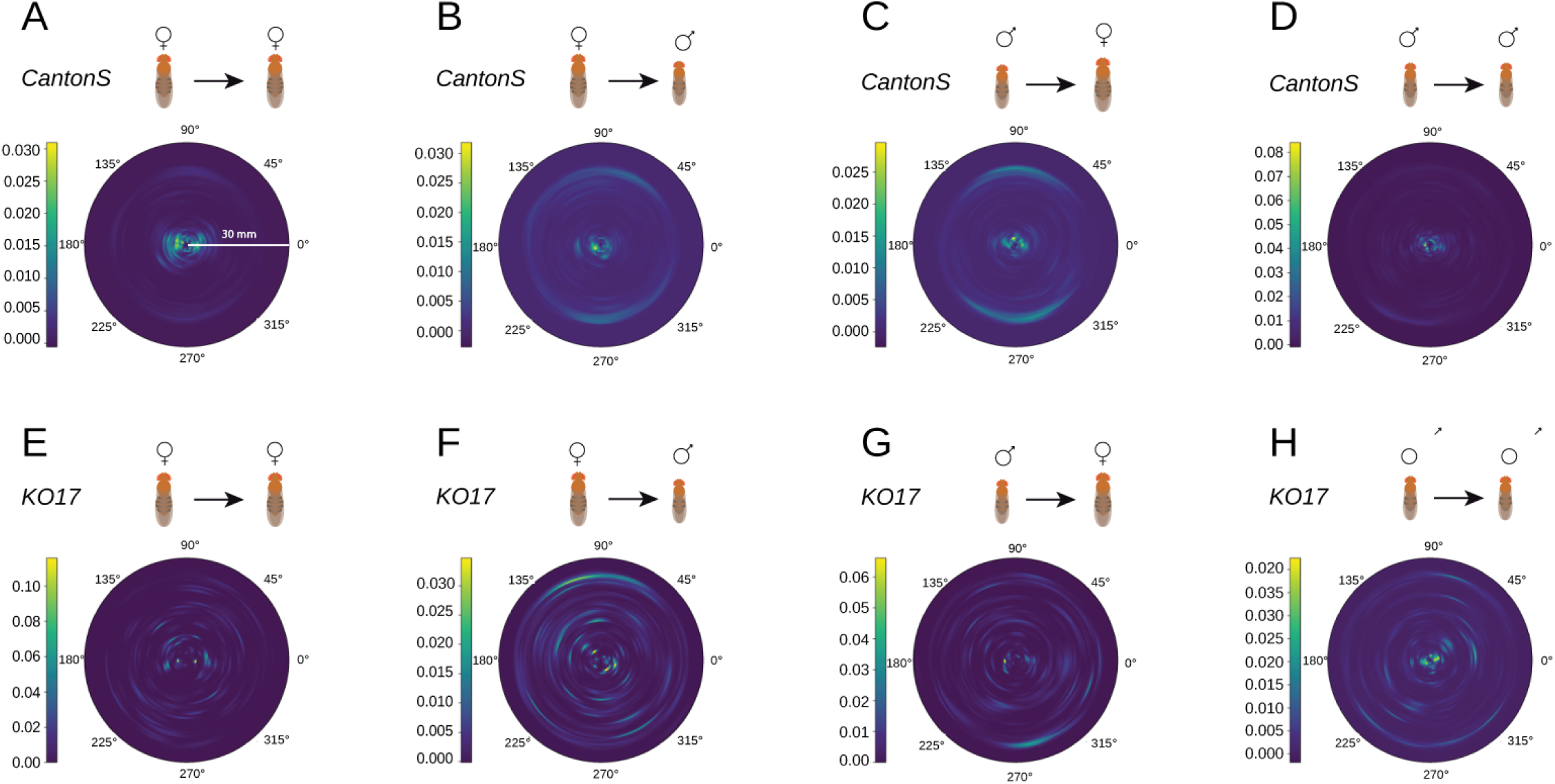
Sex-specific histograms of inter-individual distance (IID). Polar histograms are plotted as in Figure 2, with the difference that now also the sex of the focus animal is plotted separately. A) and E) show the positioning of females to each other. B) and F) show the positioning of males to female focus flies. C) and G) show the positioning of females to male focus flies. D) and H) show the positioning of males to each other. The wildtype CantonS is quantified in subplots A-D and the neuroligin-2 mutant *dnlg2*^*ko17*^ results are depicted in E-H.

A male focus fly positions male conspecifics either at a large distance of 30 mm or in close proximity < 5mm(Figure 3D). *Drosophila* male-male close interaction consists of agonistic behaviours, often in a competitive courtship setting [64, 69, 70].

If the sex of the focus fly and the conspecifics differ we observe a distribution in two distance bands. These two distances are representative of both parts of *Drosophila* courtship: the far distance and the close proximity behavious. The outer distance band ranges between 10-15 mm and is representing the far distance courtship, consisting of orientation towards the female, following behaviour and production of the male courtship song [59, 64–66, 71]. The second distance band shows the conspecifics in very close proximity of < 5 mm, oriented towards the head or the back of the focus fly and representing short distance courtship such as licking, tapping, production of courtship song and copulation [64–66].

Looking at these different constellations in the dlng mutant flies we are not able to detect a clear positioning pattern (Figure 3 E-H).

## 7 Discussion

In this study we present a new assay allowing an in-depth characterisation of social and group behaviours. Our “Social Chamber” enables the monitoring of several different modes of social behaviours and group formation, starting with a single individual and adding up to 24 animals over a predefined time period. This assay presents a low-cost and easily accessible option to assess social behaviours for small labs or in an educational setting. The recorded videos can subsequently be analysed by all commonly used tracking softwares (e.g. Lace [57]).

As a proof-of concept, we reanalysed the positional data from Corthals et. al 2017 [1] and present data on the basis of group formation, the inter-individual distance (IID). In the 2017 study the social behaviour of wildtype flies was compared to genetic knock-outs of *dnlg2* (*dnlg2*^*ko17*^). We previously calculated the nearest neighbour distance as the average distance between male individuals within already formed groups. While the nearest neighbour distance did not significantly differ between wt and *dnlg2*^*ko17*^ males within a group, the probability to be part of a group was significantly decreased in *dnlg2*^*ko17*^ males [1]. Here, we did not focus on the nearest neighbour distance of individual males, but analysed the positioning of all conspecifics in regard to the sex of the animals.

The wt fly strain *CantonS* displays a clear distribution of the specimens’ orientation, representing either courtship attempts (both at close and far distance) or male-male aggression. However, in the neuroligin mutant strain *dnlg2*^*ko17*^ no clear pattern can be observed. This is in agreement with previous findings of impaired courtship behaviour and failure to respond adequatly to surrounding accoustic signals produced by conspecifics [1, 26].

The above analysis and the 2017 study [1] illustrate the variety of parameters of social behaviour that can be quantified with an controlled social environment in insects. Social behaviour studies become more important due to the increased usage of insects as model organisms for neurological disorders [26, 72–77]. Hence it is important to establish an fundamental behavioural assay that is comparable between strains and even species.

We hope that the inter-individual distance can be used as such an fundamental parameter and that the here presented setup is an ideal tool to quantify the IID. Furthermore, observation and analysis of social behaviour can be used to teach students about the main principles of neuroscience. In an educational setting there is a need for low-cost experimental setups that can easily be produced in a larger number to allow each student firsthand experience [53].

## 8 Acknowledgements

The authors thank Martin C. Göpfert for funding, comments on the manuscript, and plenty of fruitful discussions. We are thanking Tobias Mühmer for providing the technical drawings of our arena. Many thanks to thank Alina Heukamp and Nina Hahn for extensive testing and support during the development of the setup. Special thanks go to Christian Spalthoff for providing his beautiful illustrations.

## 10 Author contribution

BG, HG and RH designed the study. SN constructed the arena with input from BG and HG. KC recorded and tracked the data which was subsequently analysed by KC and LH. The manuscript was written by BG and KC with intellectual contributions and comments from all authors.

## 11 Ethics statements

### 11.1 Studies involving animal subjects

Ethical review and approval for animal studies are not required when working on insects.

### 11.2 Studies involving human subjects

No human studies are presented in this manuscript.

### 11.3 Inclusion of identifiable human data

No potentially identifiable human images or data is presented in this study

Data availability statement Generated Statement: The original contributions presented in the study are included in the article/supplementary material, further inquiries can be directed to the corresponding author/s.

## 12 Supplementary Material

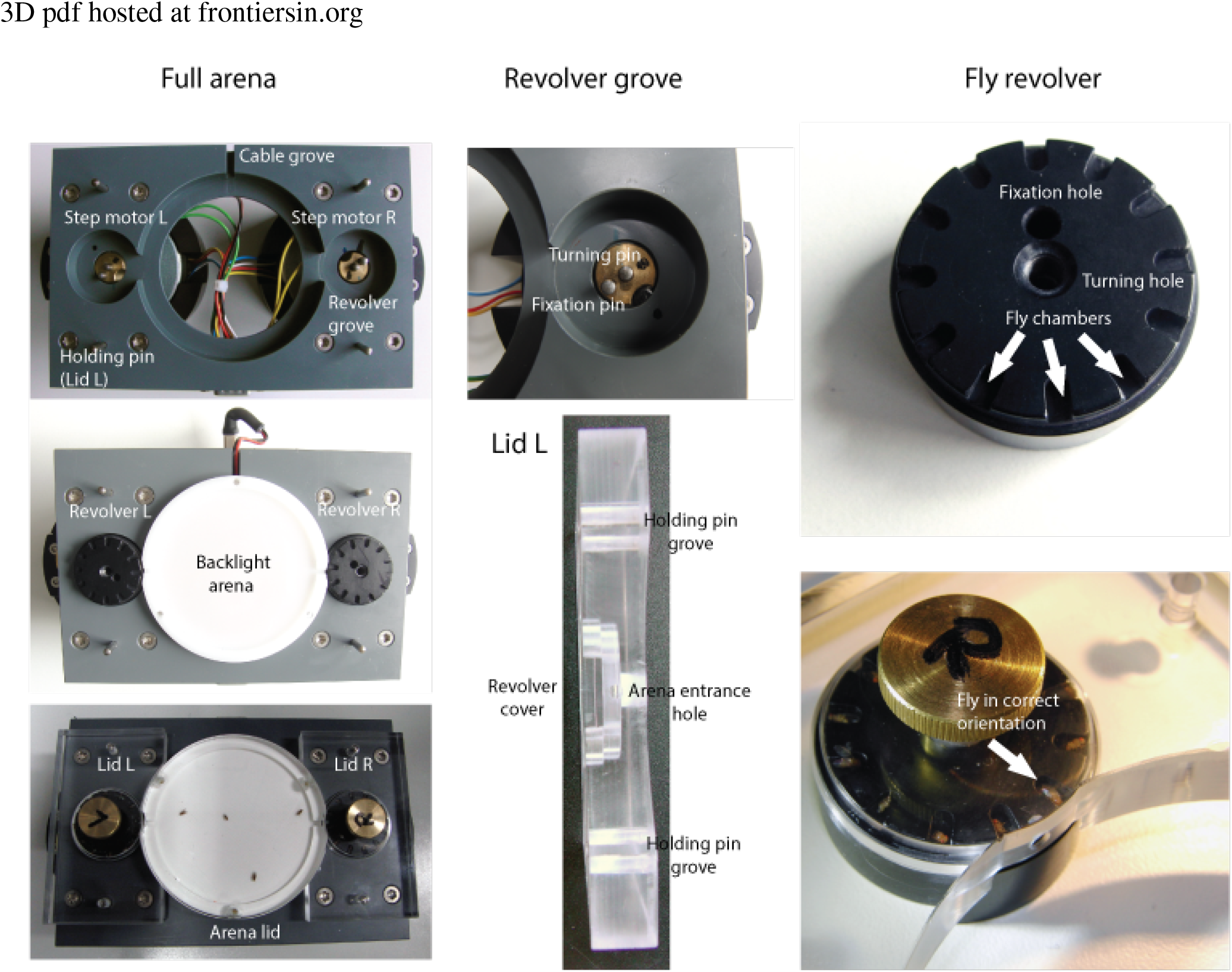

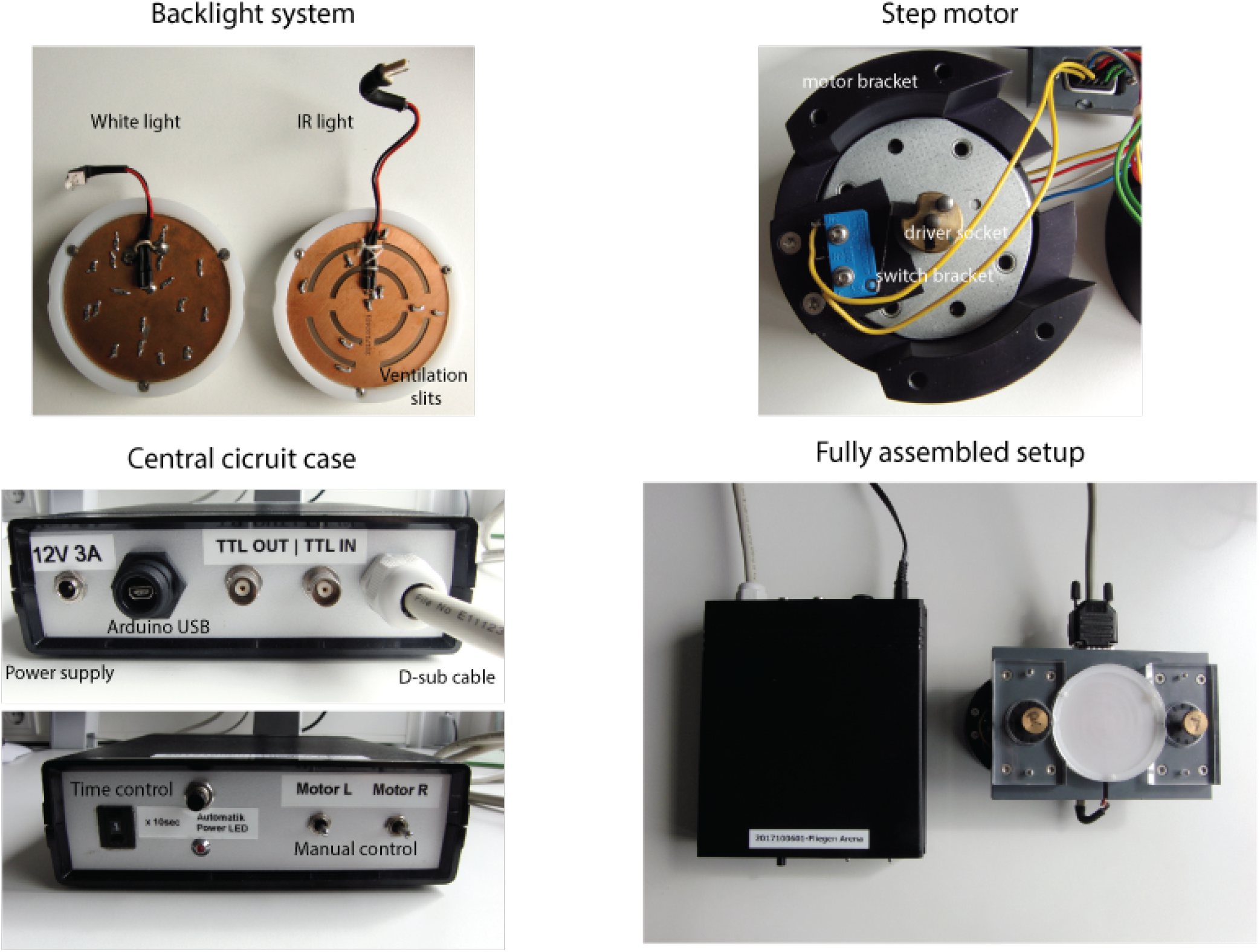

## 13 Building Plans and Circuit Diagrams

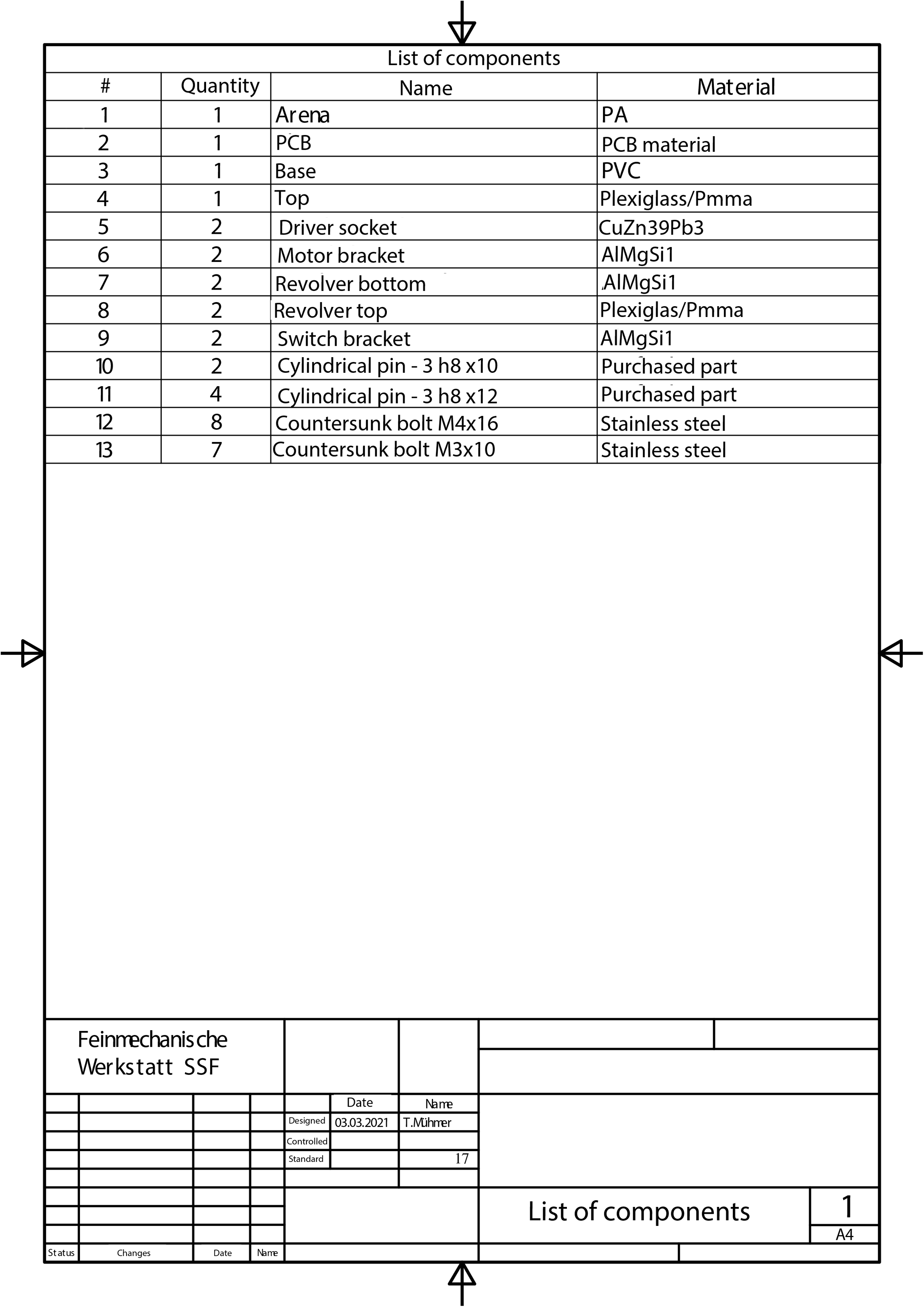

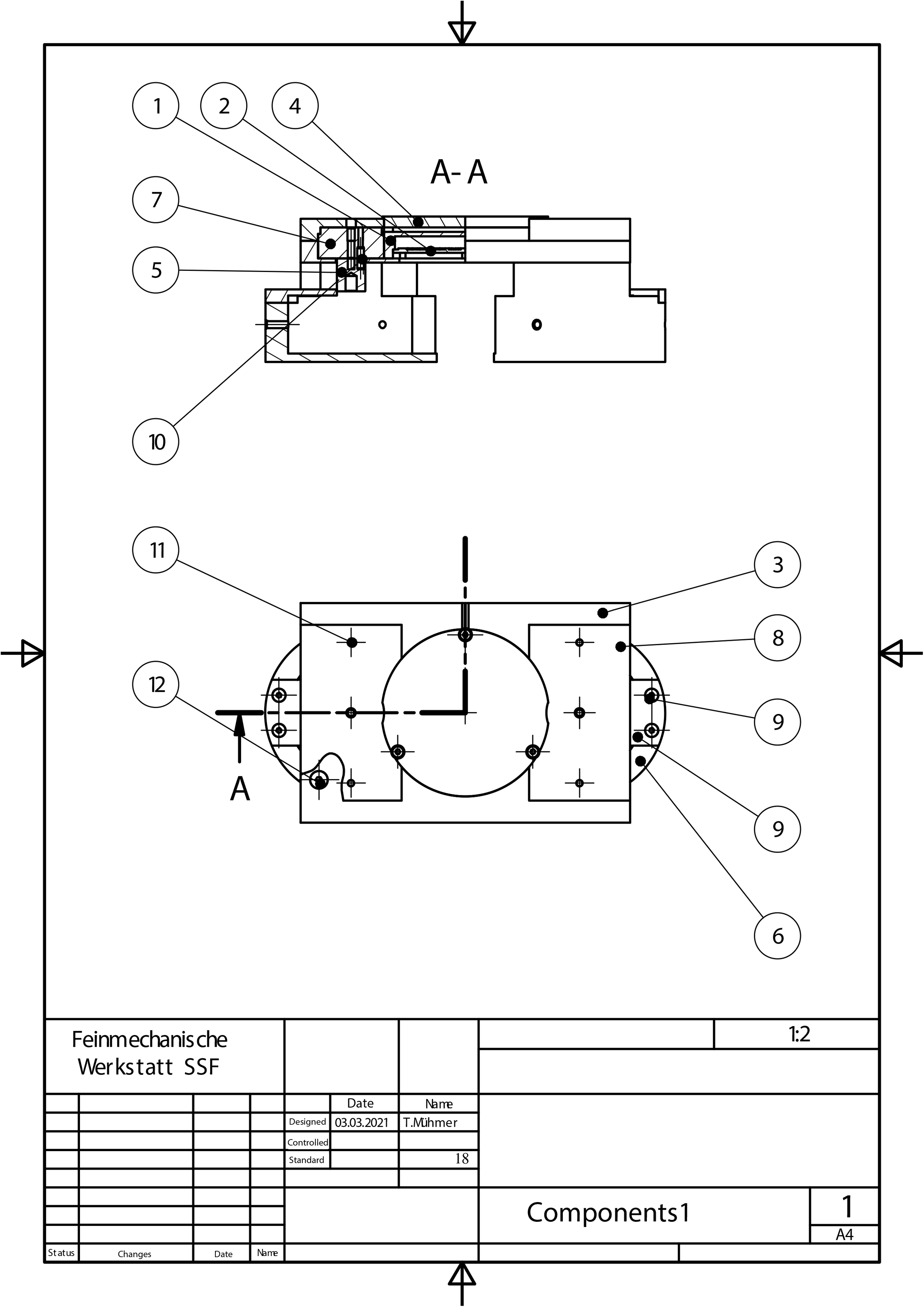

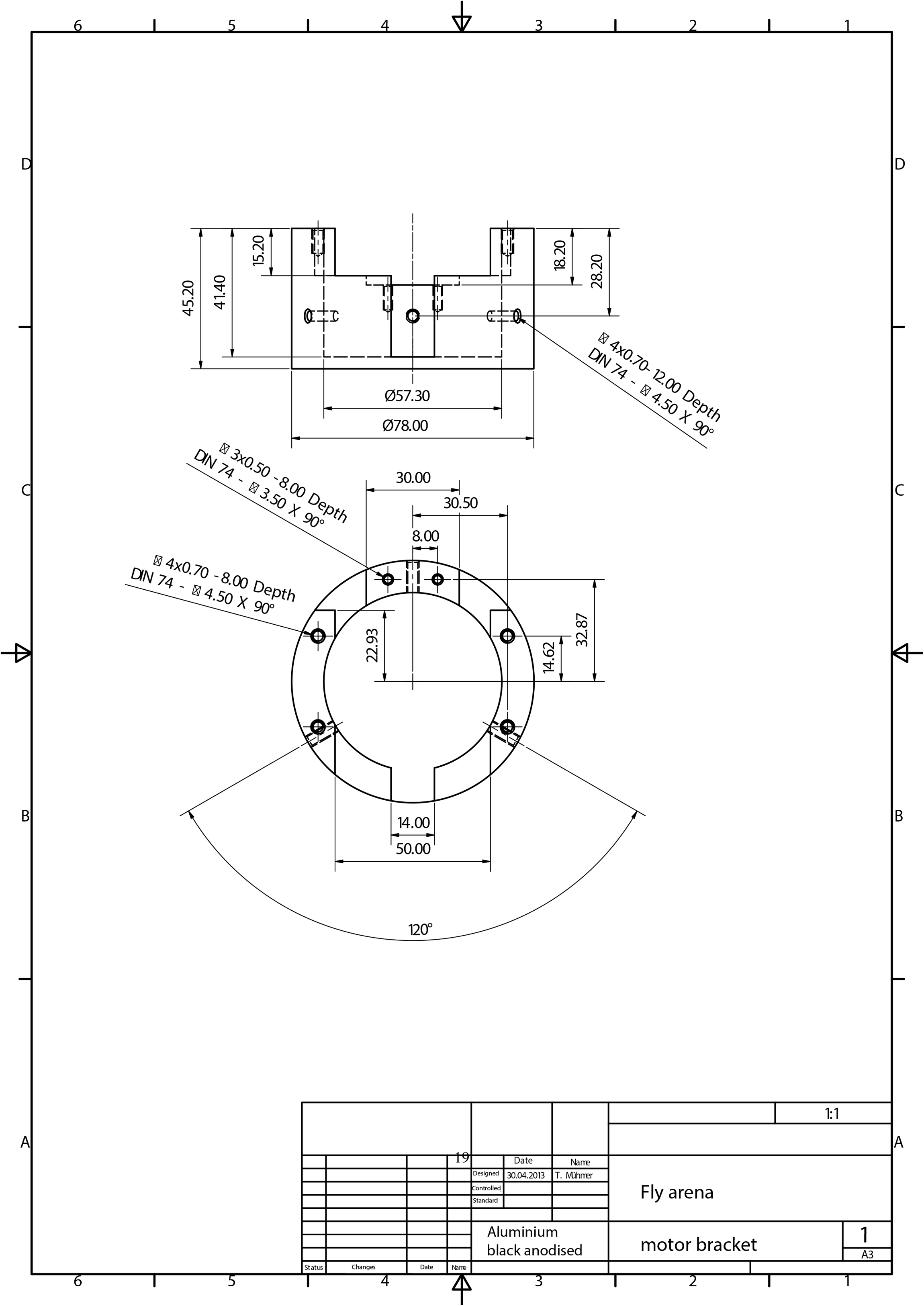

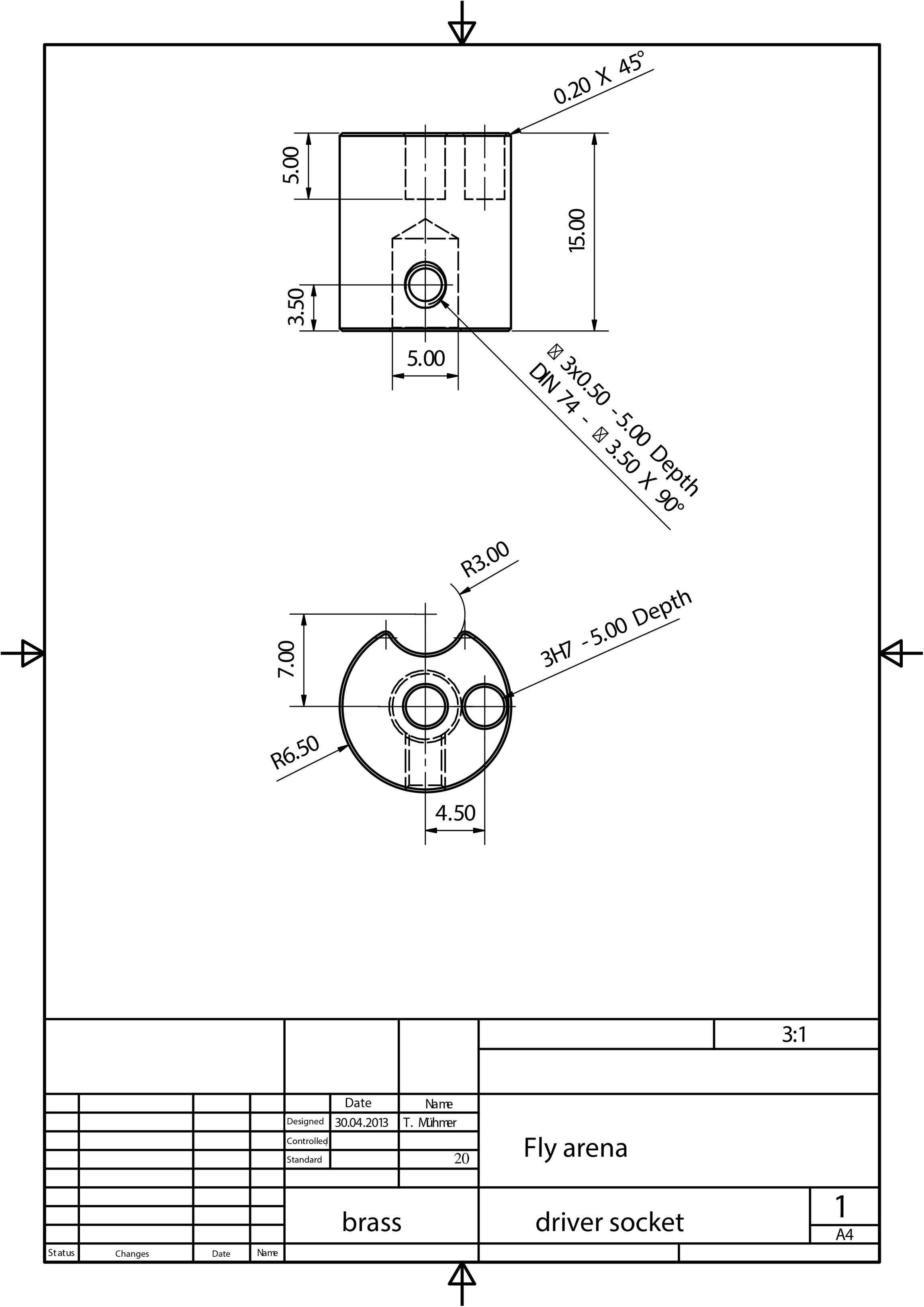

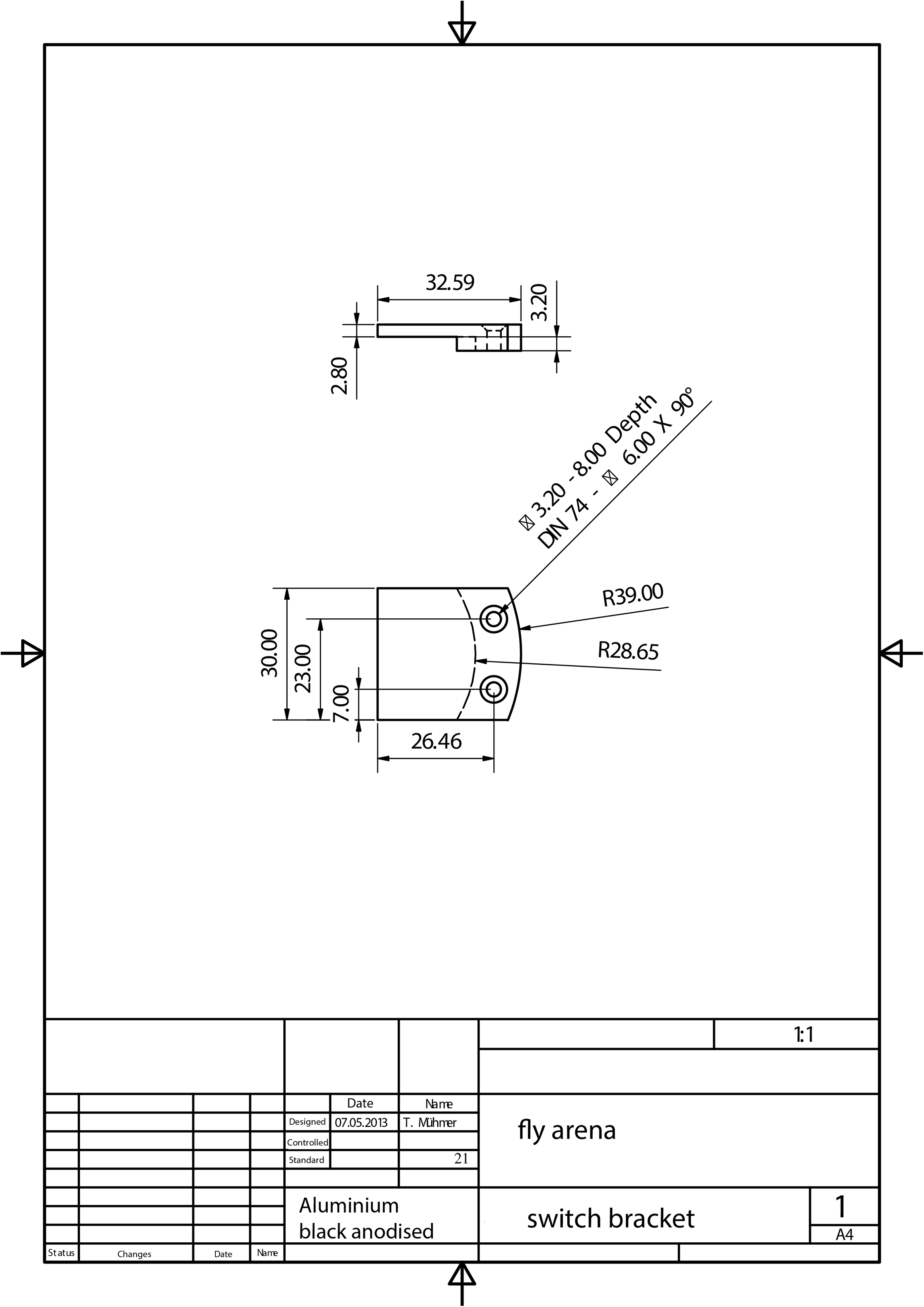

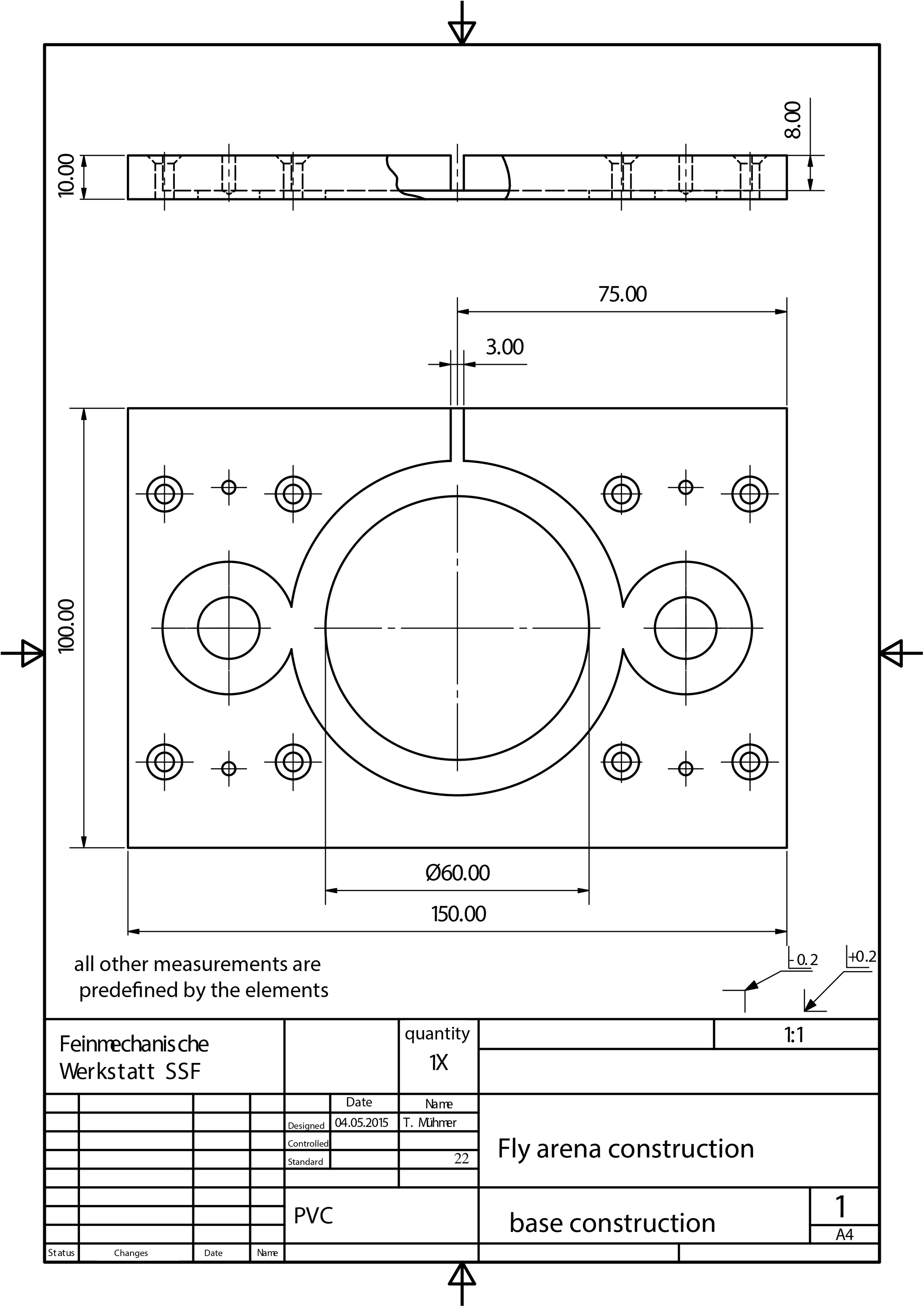

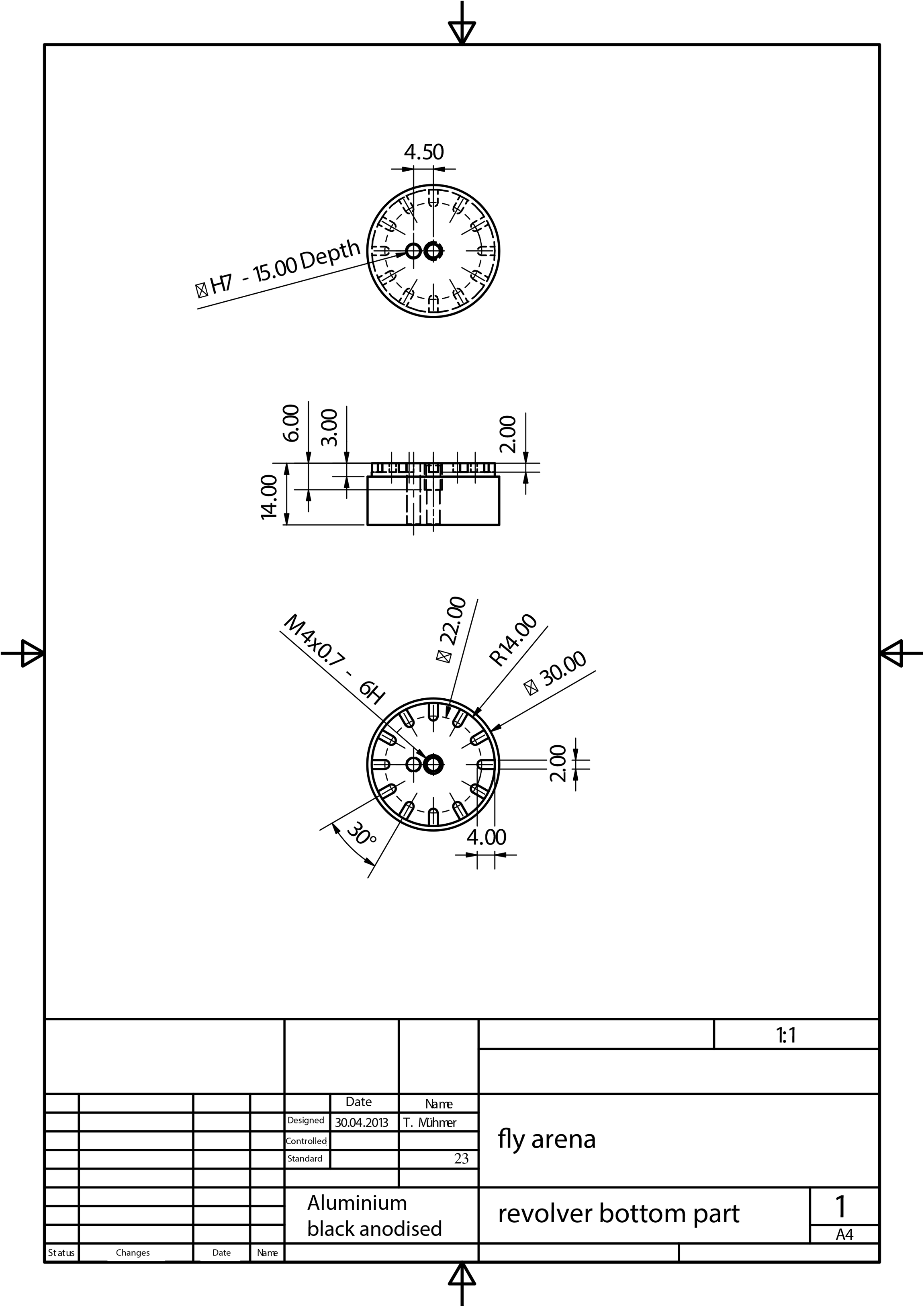

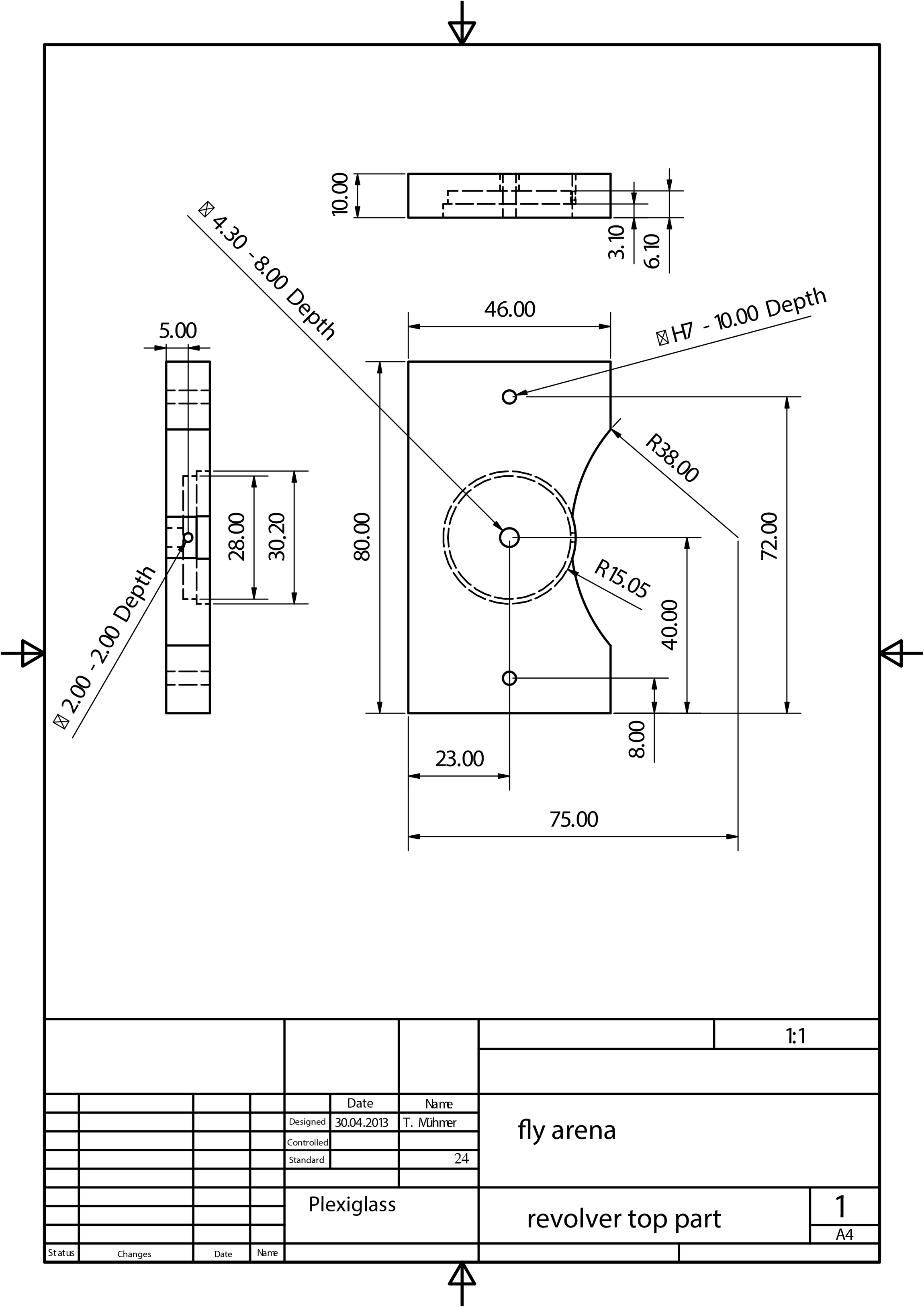

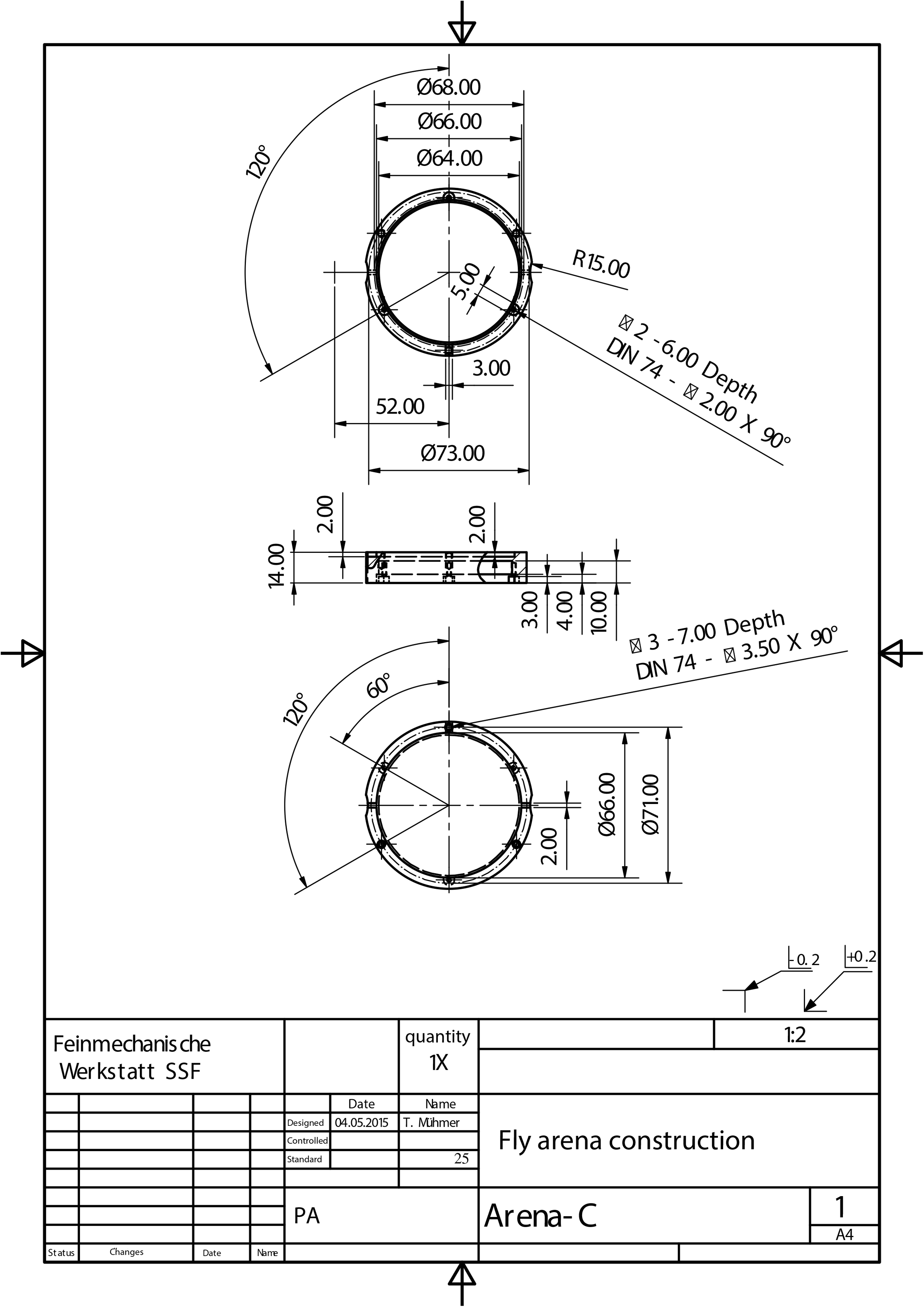

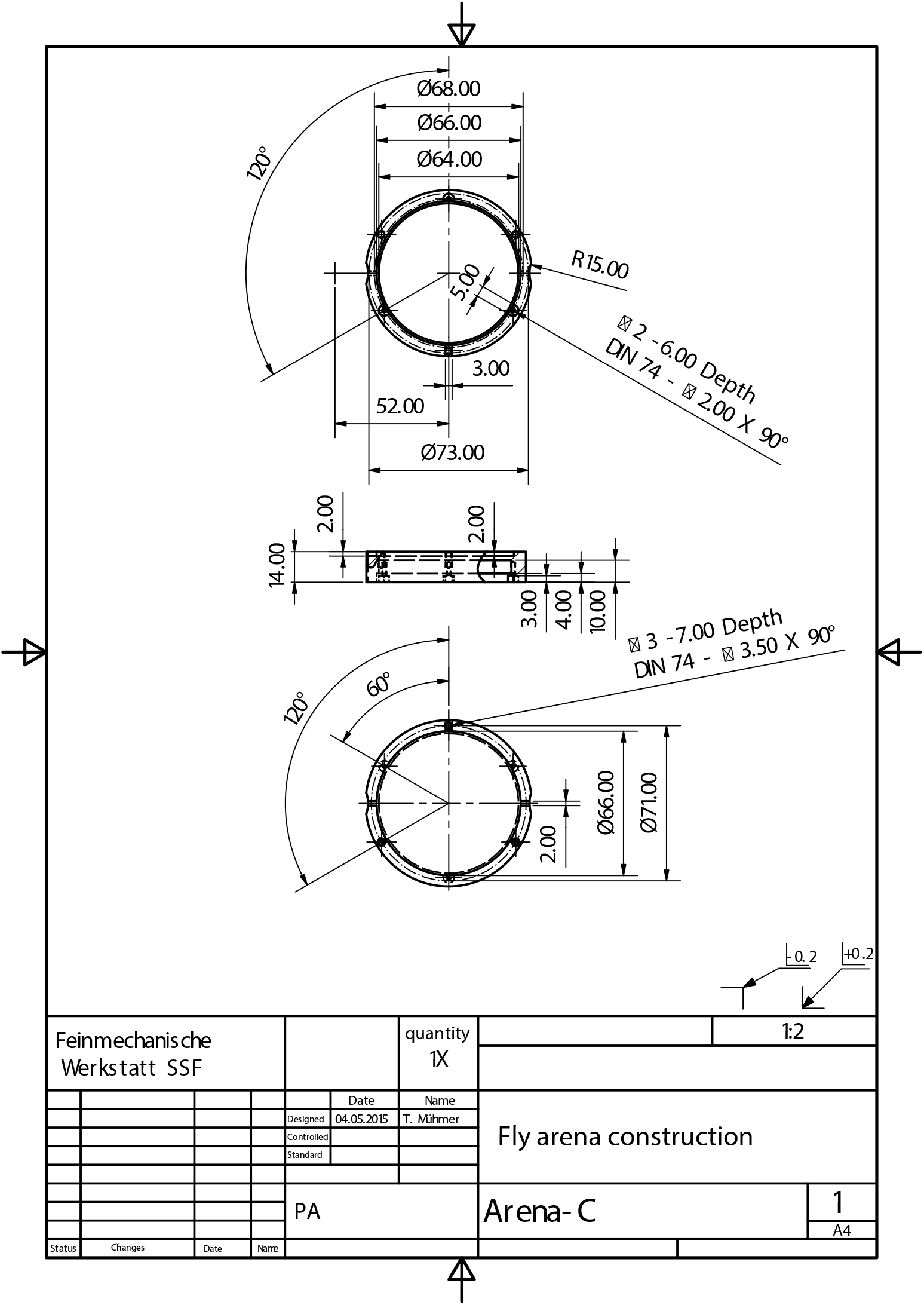

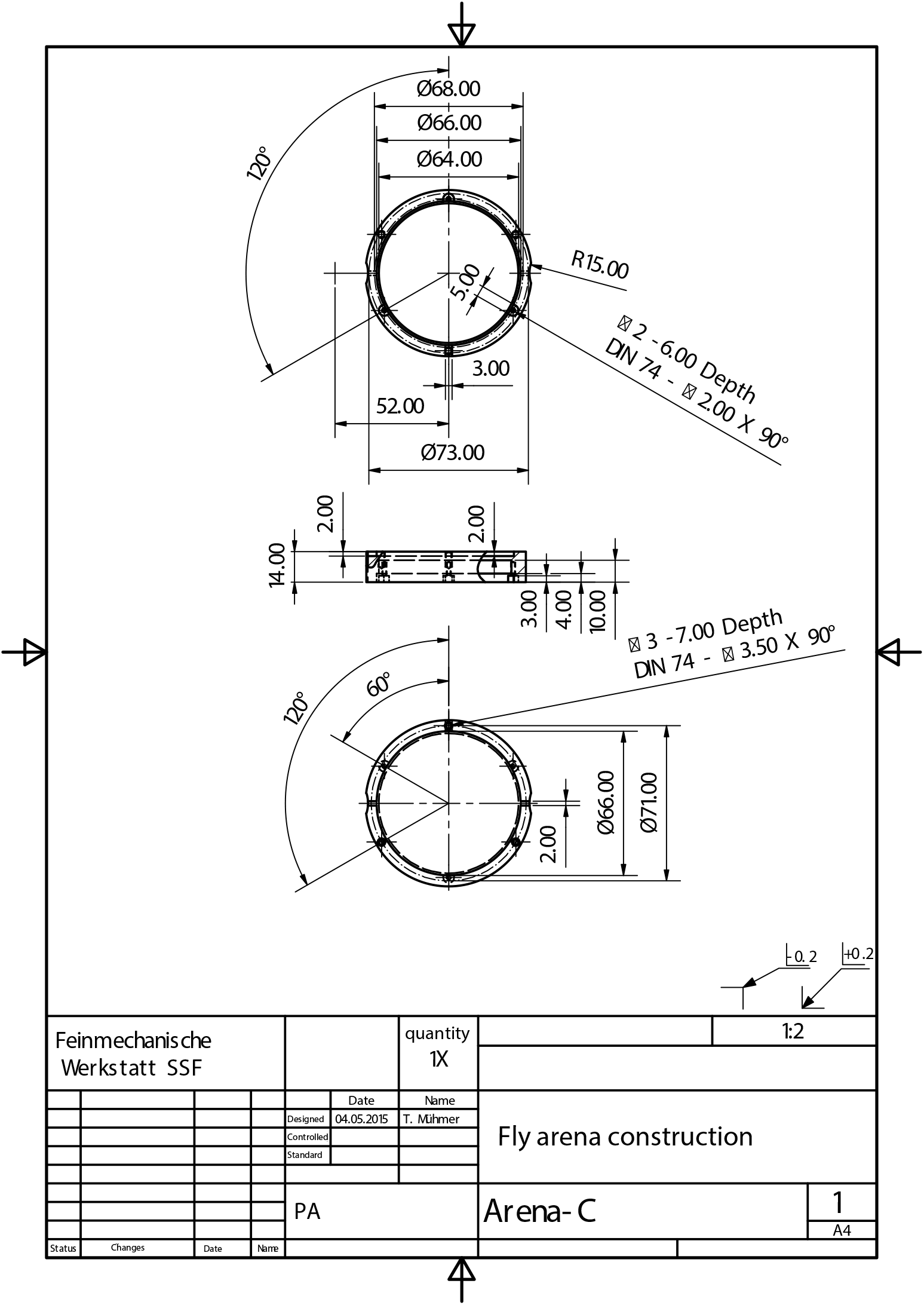

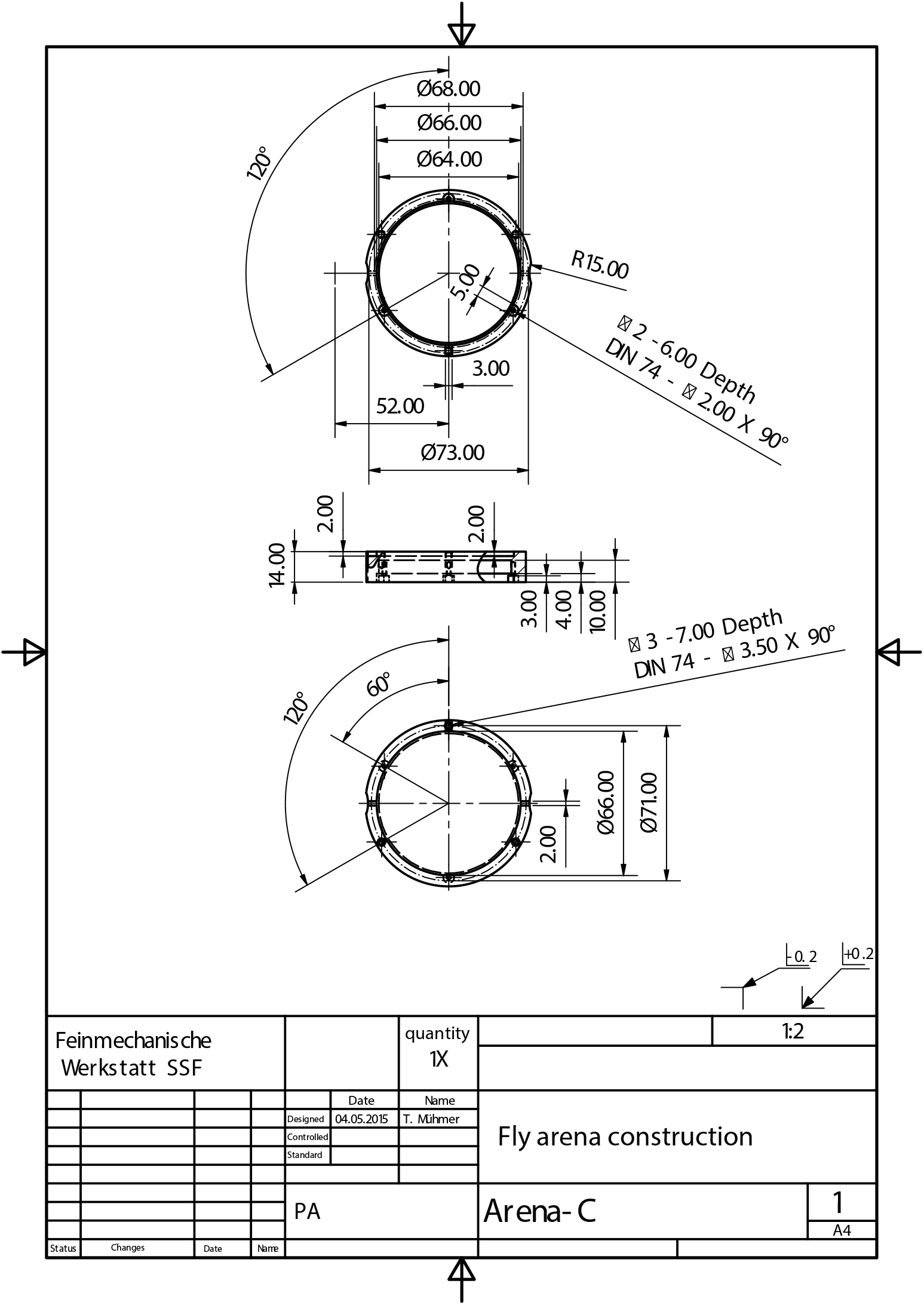

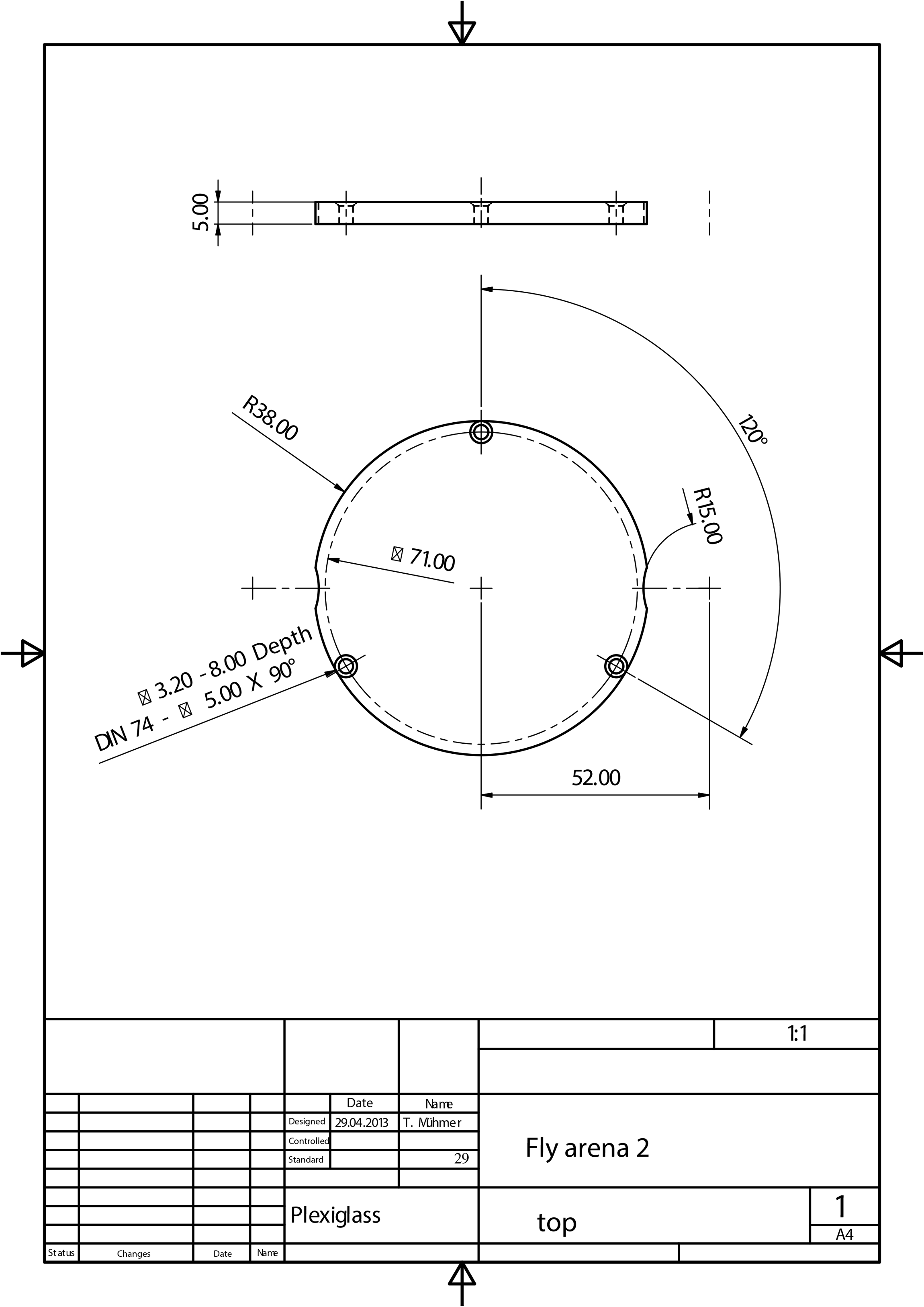

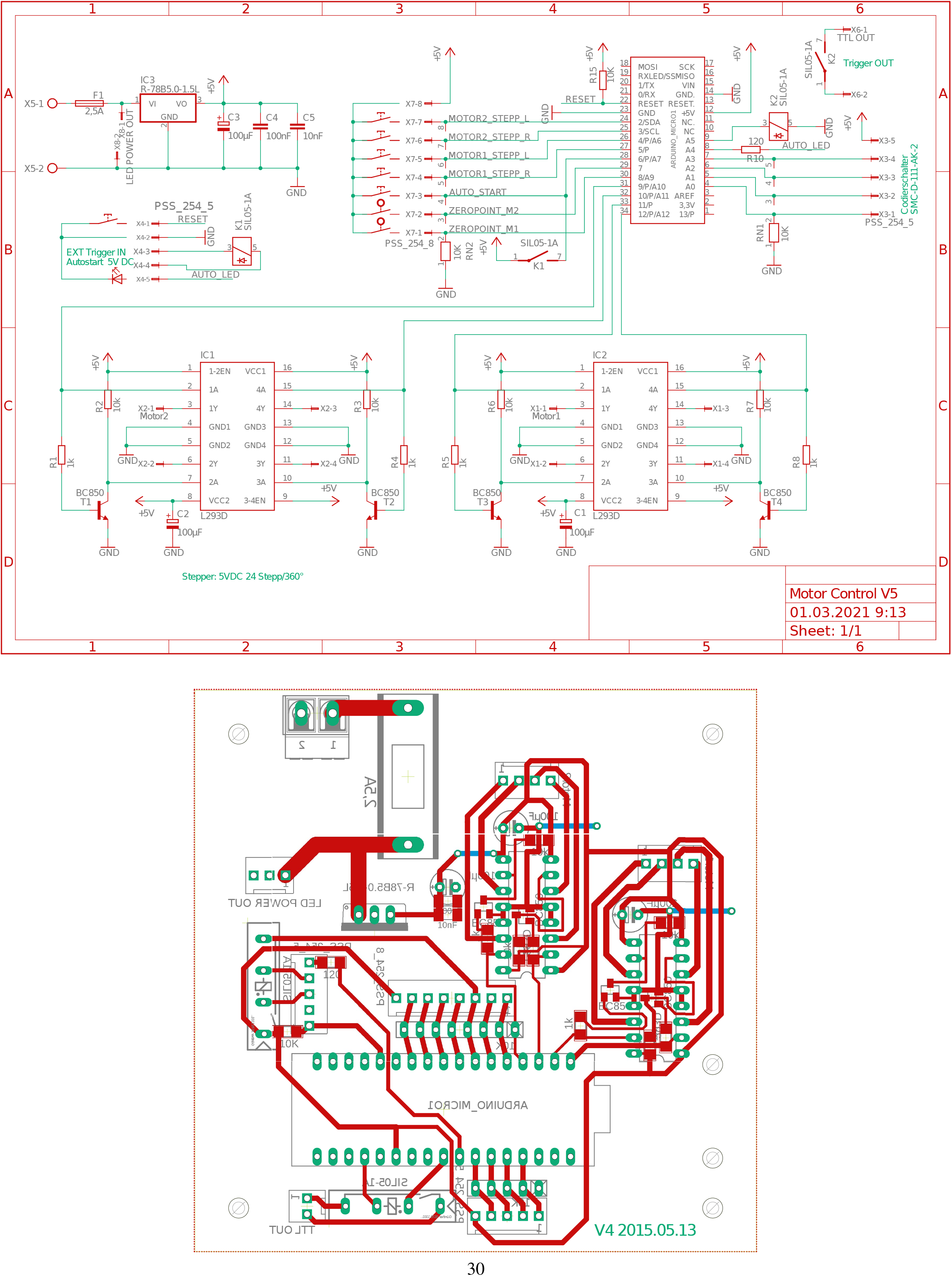

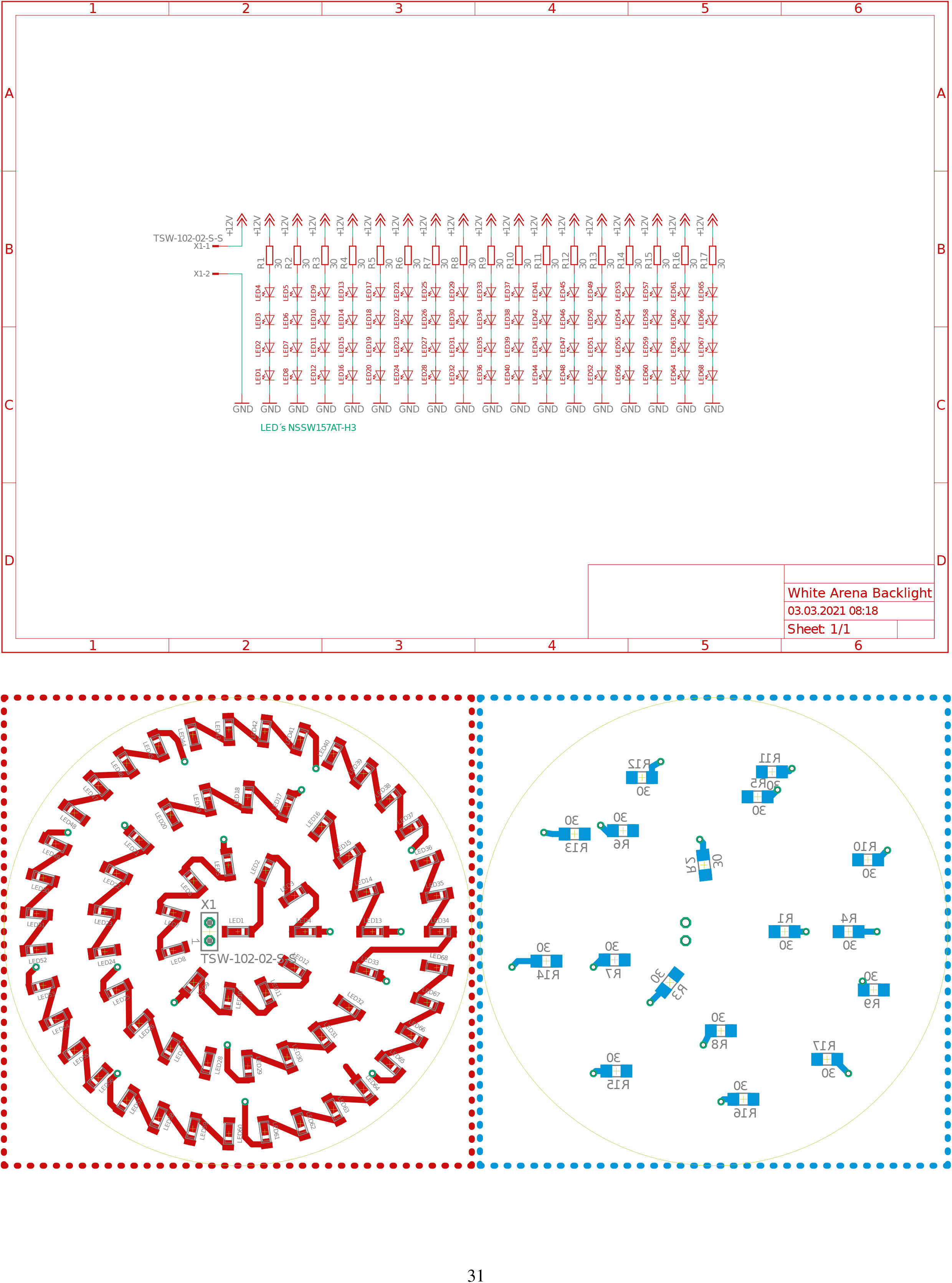

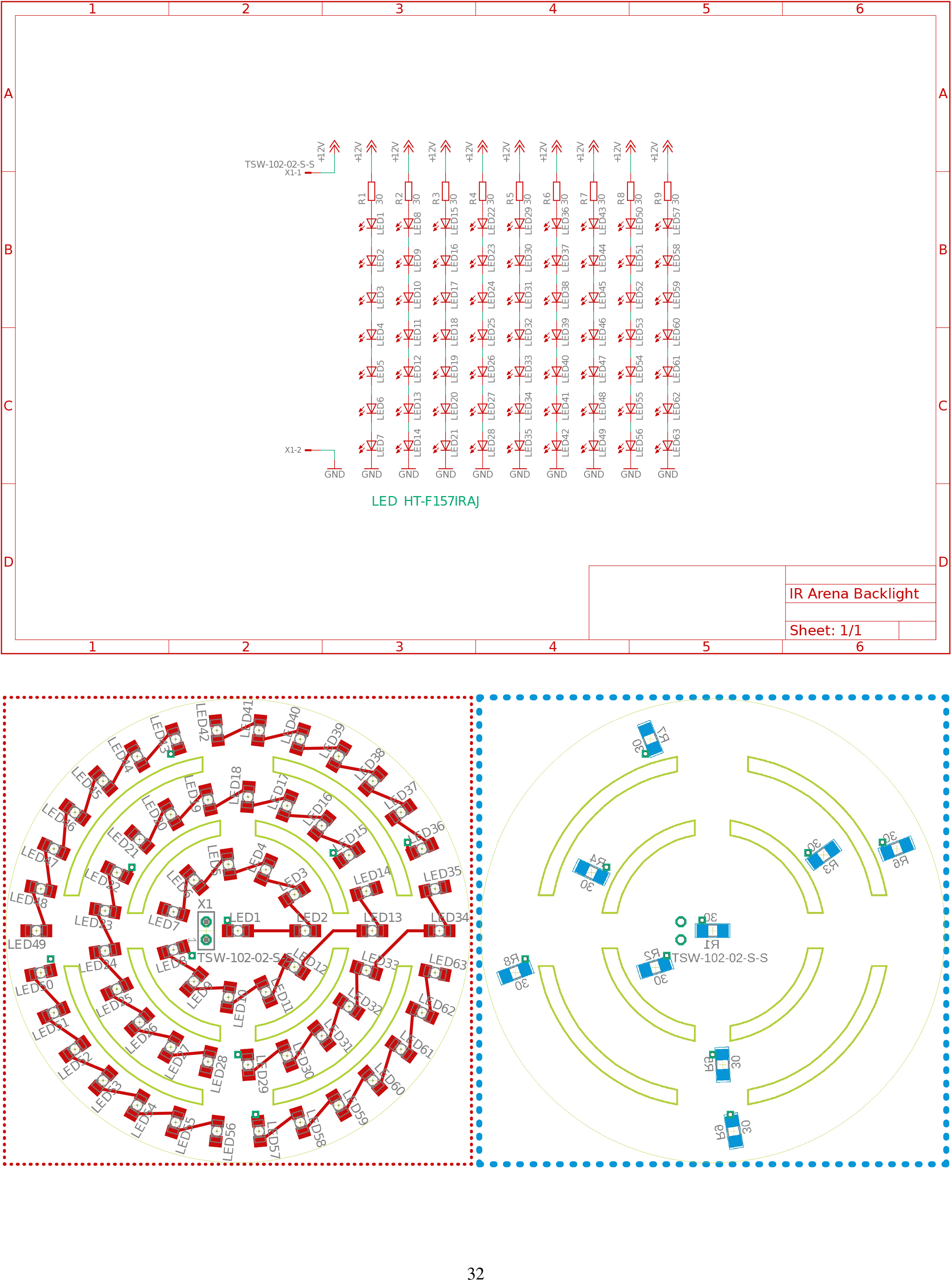

